# Human endometrial KISS1R inhibits stromal cell decidualization in a manner associated with a reduction in ESR1 levels

**DOI:** 10.1101/2022.11.20.517219

**Authors:** Jennifer Schaefer, Sangappa B. Chadchan, Ashley F. George, Nadia R. Roan, Moshmi Bhattacharya, Ramakrishna Kommagani, Andy V. Babwah

**Author notes:** **Corresponding author and person to whom reprint requests should be addressed.** Dr. Andy V. Babwah, Associate Professor, Department of Obstetrics, Gynecology and Reproductive Health, Rutgers New Jersey Medical School, Newark, NJ, USA, **Email:**.

## Abstract

Defective endometrial stromal cell decidualization is a major cause of recurrent implantation failure (RIF), a condition with a prevalence of ∼15%. To treat RIF, a stronger understanding of the endometrial factors that regulate decidualization is required. Here we studied the role of the kisspeptin receptor (KISS1R) in regulating human endometrial stromal cell (HESC) decidualization. Our data revealed KISS1R inhibits HESC decidualization in vitro in a manner associated with a striking reduction in ESR1 protein levels. To determine whether KISSR inhibition of decidualization results from reduced ESR1 levels we expressed the dominant negative ESR1-46 isoform in decidualizing HESCs. We found that expression of ESR1-46 in decidualizing HESCs ablated the expression of ESR1-66 and ESR1-54 isomers, and blocked decidualization. Interestingly, when ESR1-64 was co-expressed with ESR1-46, ESR1-66 and ESR1-54 expression was restored and decidualization was rescued. Taken together, these results suggest that KISS1R inhibits HESC decidualization by downregulating ESR1 levels. Based on our findings, we suggest that by inhibiting HESC decidualization, KISS1R regulates the depth of embryo invasion of the stroma, a requirement for a successful pregnancy.

## INTRODUCTION

Recurrent implantation failure (RIF), with an estimated prevalence of 15%, defines cases in which women have had three failed in vitro fertilization (IVF) attempts with good quality embryos (Busnelli *et al*., 2020). The underlying cause of RIF can be male (eg., sperm quality), uterine or embryo factors, or even the specific type of IVF protocol employed (Bashiri *et al*., 2018). When uterine, it may involve a uterus that is morphologically normal in appearance thus pointing to molecular defects that prevent a healthy embryo from implanting.

The human endometrium prepares for successful embryo implantation in the secretory phase by acquiring a receptive phenotype and undergoing stromal cell decidualization. Stromal cell decidualization refers to the differentiation of stromal fibroblasts into secretory epithelioid cells that contribute to the formation of the decidua, a complex tissue that additionally includes glandular epithelial cells, immune cells, and blood and lymph vessels (Radovick *et al*., 2019). The decidua regulates the depth of embryo implantation into the endometrium, selects against low-fitness embryos, and nourishes and protects the implanting embryo until the functional placenta develops (Kong *et al*., 2021; Rawlings *et al*., 2021). In the human endometrium, stromal cell decidualization begins in the mid-secretory phase in the lower portion of the functionalis surrounding the terminal spiral arteries. Decidualization then expands rapidly in a luminal-proximal direction with the functionalis becoming completely solidified with decidualized stromal cells in the mid-secretory phase (Gellersen *et al*., 2014).

Human endometrial stromal cell (HESC) decidualization is a highly regulated process and in vivo, occurs in response to rising P4 and cAMP levels in the E2-primed endometrium. Compared to P4 alone, cAMP exerts a greater effect on decidualization (Huang *et al*., 1987; Tang *et al*., 1993; Tang *et al*., 2005; Telgmann *et al*., 1998). However, when combined with P4, decidualization is enhanced (Brosens *et al*., 1999; Ferrari *et al*., 1995; Gellersen *et al*., 2003; Vinketova *et al*., 2016). Mechanistically, cAMP upregulates the expression of transcription factor FOXO1 (Takano *et al*., 2007) that induces expression of IGFBP1 (Kim *et al*., 2005) and PRL (Christian *et al*., 2002). Decidualization occurs with the increased expression of stromal ERα (Kaya Okur *et al*., 2016) and cAMP augments E2 signaling by facilitating Mediator 1 (MED1) phosphorylation. Phosphorylated MED1 binds ESR1 and gets co-recruited to estrogen response elements on ESR1 target genes where it functions as a positive co-regulator of ESR1 inducing FOXO1 and WNT4 expression (Kaya Okur, et al., 2016). The importance of ESR1 signaling as a major regulator of decidualization is further established by the observation that deletion of *Esr1* in the mouse prevents decidualization (Pawar *et al*., 2015).

Estrogen receptor alpha (ESR1) consists of multiple isoforms (**Supplemental Fig. 1**). These include full-length ESR1-66 and truncated two isoforms that include ESR1-54 and ESR1-46. ESR1-54 lacks exon 4 within the hinge region while ESR1-46 lacks the N-terminal activation function (AF-1) domain that weakens ESR1-46 transcriptional activity compared to ESR1-66 and ESR1-54 (Chantalat *et al*., 2016; Flouriot *et al*., 2000; Keselman *et al*., 2015). Studies have reported ESR1-46 acts as a dominant negative isoform by heterodimerizing with ESR1-66 to form transcriptionally weakened ESR1-66/46 heterodimers. These heterodimers then compete with the functional ESR1-66/66 homodimers for binding the estrogen response element (ERE) on target genes (Chantalat, et al., 2016; Flouriot, et al., 2000; Penot *et al*., 2005).

The kisspeptin receptor is a G protein-coupled receptor (GPCR) that is activated by a family of peptides called kisspeptins (KPs). KISS1R and KP are expressed in the pregnant mouse uterus, and we have previously demonstrated that KISS1R regulates early mouse pregnancy by negatively regulating epithelial ESR1 mRNA and protein expression and transcriptional activity at the time of implantation (Calder *et al*., 2014; Fayazi *et al*., 2015; Kotani *et al*., 2001; León *et al*., 2016; Pampillo *et al*., 2009; Schaefer *et al*., 2021; Szereszewski *et al*., 2010). KISS1R and KP are also expressed in the human endometrium (Baba *et al*., 2015; Cejudo Roman *et al*., 2012; Hugon-Rodin *et al*., 2018). While their functions in the endometrium are not well understood, it was shown that human endometrial stromal cells (HESCs) subjected to in vitro decidualization for 12 days resulted in a small increase in *KISS1* but not *KISS1R* mRNA levels, suggesting a possible role for KP in regulating HESC decidualization. However, this role was never directly addressed.

The goal of the current study is to further characterize the expression of KISS1R and KP in the healthy human endometrium during the secretory phase and determine their function. Based on our findings, we report that KP/KISS1R signaling inhibits HESC decidualization in a manner associated with the downregulation of ESR1 expression. Based on our data, we suggest that KISS1R inhibits HESC decidualization and thereby regulates the depth of embryo invasion of the stroma, a requirement for a successful pregnancy.

## MATERIAL AND METHODS

### Endometrial biopsy collection

Human endometrial samples were obtained by Pipelle biopsies in accordance with the Institutional Review Board (IRB) approved written consents from (1) Department of Obstetrics, Gynecology and Reproductive Health, Rutgers New Jersey Medical School, Newark, NJ (2) Department of Obstetrics and Gynecology, Washington University, St. Louis, MO and **(3)** Department of Obstetrics, Gynecology and Reproductive Sciences, University of California, San Francisco, CA. Endometrial biopsies were collected from the proliferative (menstrual day 9, 11, 12), early-secretory and late-secretory phases from healthy non-pregnant women (based on a negative urine pregnancy test) of reproductive age. These women exhibited regular menstrual cycles (25-32 days), had not used any hormonal/intrauterine contraceptive devices for 30 days prior to collection, were not on antibiotic treatment and did not show evidence of current vaginal infection, sexually transmitted diseases and abnormal Pap smear.

### Isolation and culture of primary HESCs

Human endometrial stromal cells were isolated from the endometrial biopsies as described (Michalski *et al*., 2018) and (Chen *et al*., 2015). HESCs were cultured at 37°C under 5% CO_2_ in phenol red-free DMEM/F-12 medium containing 10% final volume of charcoal-stripped fetal bovine serum (ThermoFisher Scientific Cat # A3382101) and penicillin and streptomycin (Michalski, et al., 2018); hereafter, this medium is referred to as basal medium (BM).

### HESC *in vitro* decidualization

Primary HESCs were induced to undergo decidualization in phenol red-free DMEM/F-12 medium containing 2% charcoal-stripped FBS, 0.01 μM estradiol (E2), 1 μM medroxyprogesterone acetate and 500 μM N6,2’-O-dibutyryladenosine 3’:5’-cyclic monophosphate (Sigma Cat # D0627); hereafter, this medium is referred to as EPC medium. Cells were subjected to the decidualization protocol for one to six days. Decidualization was confirmed visually by morphological changes from fibroblastic to epithelioid cells and increased expression of decidualization markers, including *IGFBP1* and *PRL* (Michalski, et al., 2018).

### Confocal immunofluorescence analysis

Human endometrial tissue from early- and late-secretory phases were fixed in 10% (w/v) formalin, paraffin embedded and sectioned at 5 μm. Cultured HESCs were fixed with 4% paraformaldehyde pH 7.4. Tissue sections and cells were incubated with the following primary antibodies. For tissue sections we used KISS1 rabbit polyclonal antiserum antibody (Cat # AC566, 1:10,000) (Clarkson *et al*., 2009) and KISS1R rabbit polyclonal antibody (ABCAM, Cat # ab137483, 1:100). For cultured HESCs, we used CK7 rabbit monoclonal antibody (ABCAM, Cat # ab181598, 1:2000). Slides (with tissue or cells) were then incubated with goat anti-rabbit IgG secondary antibody, poly HRP conjugate. Immunoreactivity was detected using the Alexa Fluor 555 Tyramide SuperBoost Kit with all slides processed in parallel (ThermoFisher Scientific, Cat #B40923). Stained tissue or cells were overlaid with Fluoromount-G with DAPI mounting medium (ThermoFisher Scientific, Cat # 00-4959-52) and sealed with a coverslip. A matched negative staining control, normal rabbit IgG polyclonal antibody (Sigma Aldrich, Cat # 12-370, 1:100), was used to confirm the primary antibody staining and binding specificity. Immunostained slides were analyzed by confocal microscopy.

### RNA isolation and quantitative PCR (qPCR) analysis

Gene expression was determined on total RNA prepared from non-decidualized and decidualized cells. RNA was isolated from cells using TRIzol reagent (ThermoFisher Scientific; Cat # 15596018). cDNA was synthesized using iScript™ Reverse Transcription Supermix (Bio-Rad, Cat # 1708841). Briefly, one μg of total RNA was reverse transcribed and diluted 2-fold in nuclease free water. Five μl of diluted cDNA was used in each qPCR reaction. qPCR was conducted using the iTaq Universal SYBR Green Supermix (Bio-Rad, Cat # 1725124). qPCR was performed in duplicate for each sample and done a total of three independent times using primers for the following genes **(Table 1)**: ribosomal RNA (*18SN5*), decorin (*DCN*), *DESMIN*, estrogen receptor alpha (*ESR1*), forkhead box protein O1 (*FOXO1*), insulin-like growth factor-binding protein 1 (*IGFBP1*), kisspeptin (*KISS1*), kisspeptin receptor (*KISS1R*), cytokeratin 7 (*KRT7*), cytokeratin 18 (*KRT18*), left-right determination factor 2 (*LEFTY2*), matrix metallopeptidase 2 (*MMP2*), matrix metallopeptidase 7 (*MMP7*), progesterone receptor (*PGR*), prolactin (*PRL*), and wingless-type MMTV integration site family member 4 (*WNT4*). Gene expression was calculated as relative expression to the housekeeping gene (*18SN5*), using the 2^−ΔΔCt^ method.

**Table 1.**
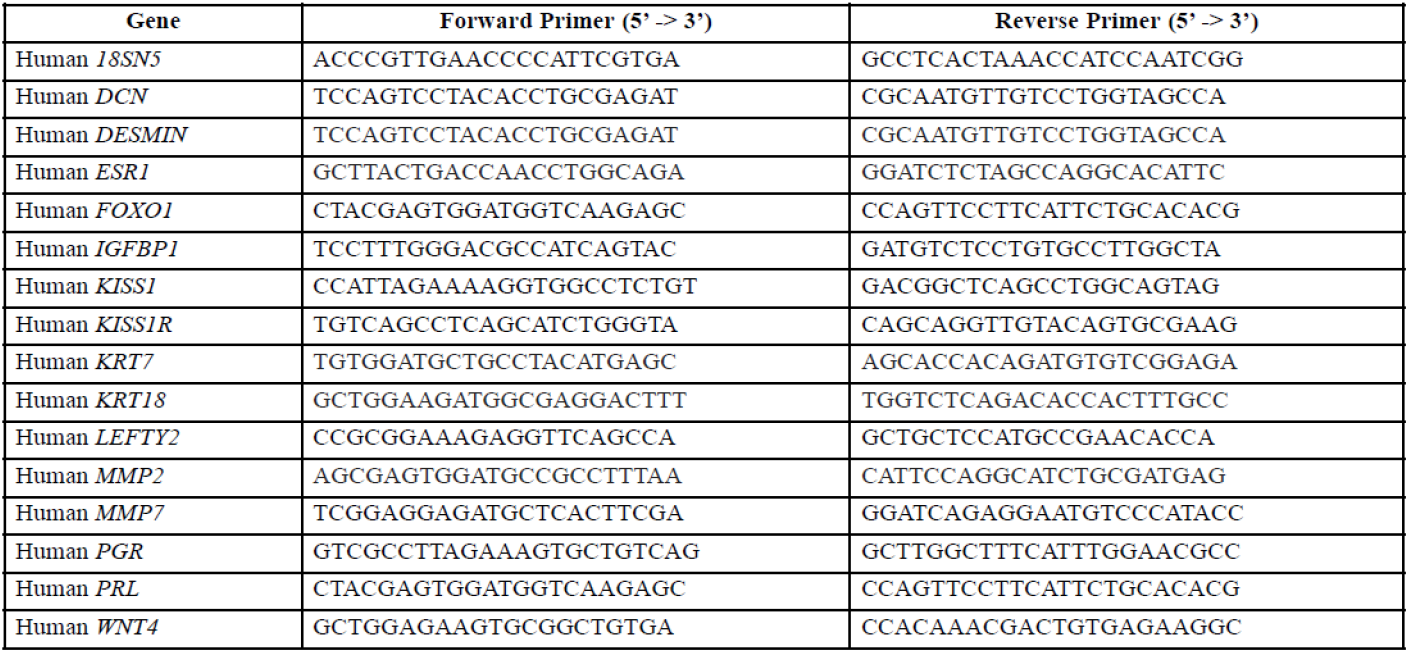
Human primer sequences used in qPCR analyses

For some studies involving primary HESCs, qPCR was also conducted using TaqMan Assays (Applied Biosystems/Life Technologies). One μg of RNA was reverse transcribed with the High-Capacity cDNA Reverse Transcription Kit (Thermo Scientific, Waltham, MA, USA). The amplified cDNA was diluted to 10 ng/μl, and quantitative PCR was performed using 40 ng of cDNA with the following TaqMan probes (Applied Biosystems/Life Technologies), *KISS1* (Hs00560041_m1), and *KISS1R* (Hs00173906_m1). Universal TaqMan 2X Mastermix (Applied Biosystems/Life Technologies, Grand Island, NY) on a 7500 Fast Real-time PCR system (Applied Biosystems/Life Technologies). Ribosomal RNA (18S) (Thermo Scientific, Waltham, MA, USA, Cat # 4318839) was used as an internal control to calculate relative expression using the 2^−ΔΔ^Ct method.

### Transfection and expression of HESCs

Primary HESCs were induced to undergo decidualization and on the fourth day of decidualization, using Lipofectamine 3000 (ThermoFisher Scientific, Cat # L3000-015), HESCs were transfected with an empty expression plasmid (control) or human *KISS1R* expression plasmid corresponding to the open reading frame (Origene) or human *ESR1* expression plasmids h*ESR1*-46 or h*ESR1*-66 (Flouriot, et al., 2000; Penot, et al., 2005). On the day of transfection, cells were treated with 100 nM KPA, a stable KP analog (Guzman *et al*., 2022). HESCs were cultured in decidualization medium for two additional days (total of six days of decidualization). On each of the two days, the KPA-supplemented decidualization medium was refreshed (Guzman, et al., 2022). In addition, HESCs were also grown in modified basal medium containing 2% charcoal stripped FBS to serve as matched negative controls for each transfection group. RNA and protein were then isolated and subjected to qPCR analysis of gene expression and western blotting for protein expression, respectively.

### Protein extraction and western blot analysis

Primary HESCs were lysed and 25 μg of cell lysates were used for western blot analysis. Proteins were separated on a 10% SDS-PAGE gel and transferred to a 0.45 μM nitrocellulose membrane. After blocking, membranes were incubated overnight with the following primary antibodies: ESR1 rabbit polyclonal antibody (Santa Cruz Biotechnology (HC-20), Cat# sc-543, 1:1000), FOXO1 mouse monoclonal antibody (Santa Cruz Biotechnology (C-9), Cat# sc-374427, 1:500), and tubulin mouse monoclonal antibody (ABCAM, Cat# ab184613, 1:5000). Subsequently, protein blots were incubated with the following secondary antibodies Anti-Mouse IgG HRP Linked Antibody (Cell Signaling, Cat # 7076S, 1:2500) and Anti-Rabbit IgG HRP Linked Antibody (Cell Signaling, Cat # 7074S, 1:2500) and visualized by chemiluminescence with ChemiDoc Touch Imaging System (Bio-Rad Laboratories) using an SuperSignal and West Dura Extended Duration Substrate ECL detection system (Thermo Fisher Scientific, Cat # 34076). Densitometric analysis of protein levels were performed using the Image Lab 6 Software (Bio-Rad Laboratories).

### Statistics

The differences between groups were determined using unpaired, two-tailed Student’s t-test or one-way ANOVA followed by post hoc Bonferroni’s Multiple Comparison test (GraphPad Prism Software, Inc, La Jolla, CA). All values are expressed as mean ± SEM and a value of *P*<0.05 was considered statistically significant.

## RESULTS

### KP and KISS1R are strongly expressed in the secretory phase human endometrium

In order to assess the expression of KP, KISS1R, and ESR1 protein, immunofluorescence was performed on secretory phase human endometrium using validated antibodies (Blake *et al*., 2017; Cvetkovic *et al*., 2013; Desroziers *et al*., 2010). In the early-secretory phase, KP was strongly expressed throughout the endometrial stroma (**Fig. 1A**) and localized to the lumen of some glands (**Fig. 1A**, see arrowheads), while KISS1R was weakly but clearly detected within the epithelium of some glands (**Fig. 1B**). ESR1 was strongly expressed in the nuclei within the glandular epithelium and moderately throughout the stroma (**Fig. 1C**). At the start of the mid-secretory phase, within the window of implantation, KP became focused around the glandular epithelium (designated the sub-glandular region) in the stroma (**Fig. 1E** and **I**) and highly localized within the glandular lumen (**Fig. 1E** and **I**, see arrowheads). In both the early- and mid-secretory phase endometrium, KP was never detected on glandular epithelial cells (**Fig. 1A, E** and **I**). In the mid-secretory phase, KISS1R was highly localized on all glands within the glandular epithelium along the basal lamina as well as throughout the stroma (**Fig. 1F** and **J**). In the mid-secretory phase, ESR1 expression was moderately localized to the glandular epithelium, stroma and cytoplasm (**Fig. 1G** and **K**, see arrowheads). Negative staining controls using an isotype equivalent IgG confirmed that the KP, KISS1R, and ESR1 staining was specific (**Fig. 1D, H**, and **L**). Collectively, these results demonstrate that based on the spatial distribution of KP and KISS1R, stromal KP likely activates stromal and epithelial KISS1R which could then regulate ESR1 expression and signaling.

**Fig. 1.**
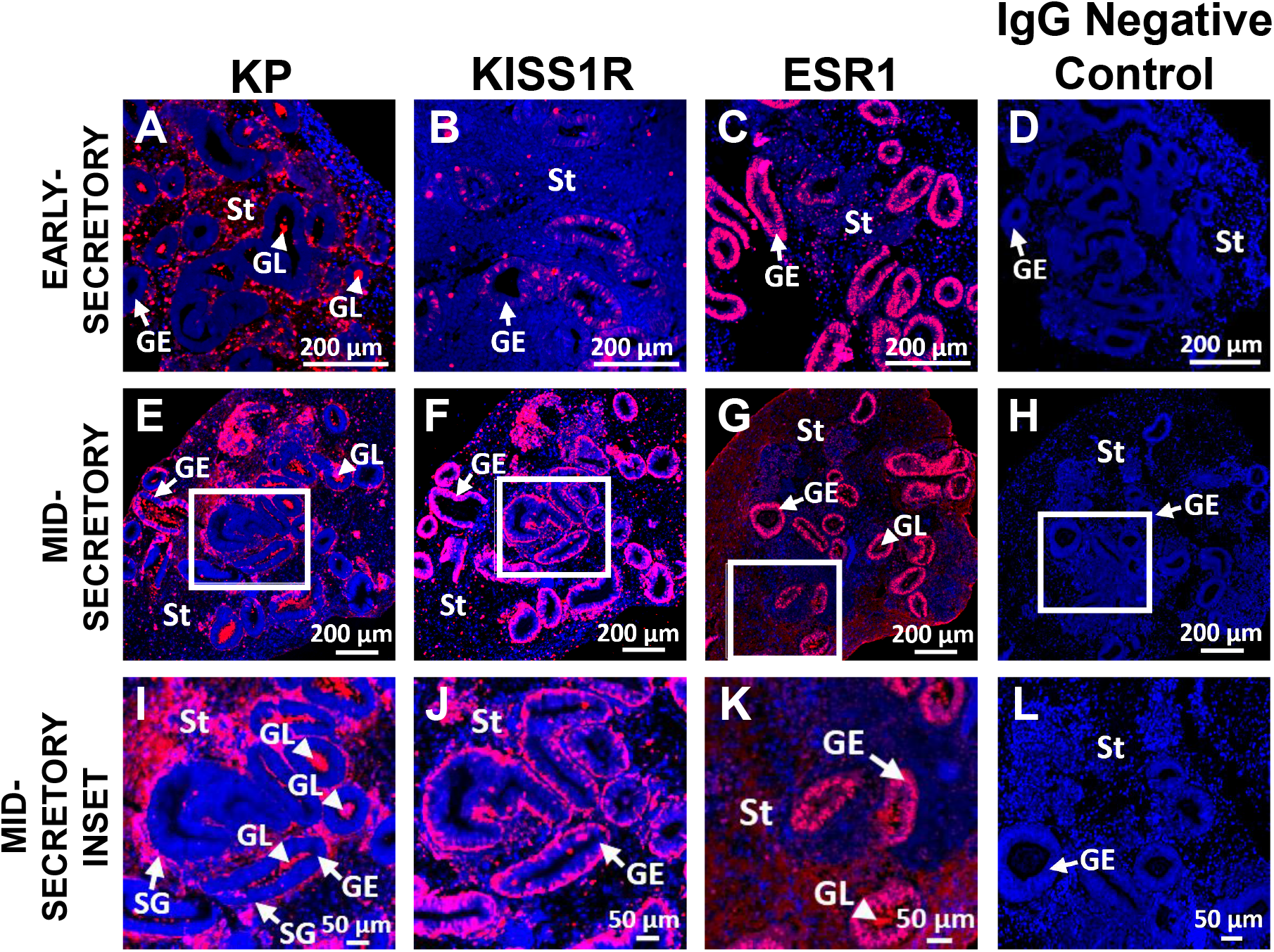
KP and KISS1R are strongly expressed in the endometrium in the mid-secretory endometrium. Immunofluorescence analysis of KP (A, E and I), KISS1R (B, F and J), and ESR1 (C, G and K) protein in the human endometrium during the early- and mid-secretory phase of the menstrual cycle. I, J, K and L are magnified sections shown in E, F, G, and H respectively. D, H and L are negative staining controls using an IgG antibody to confirm that the primary antibody staining in the endometrium is specific. Arrows point to the glandular epithelium (GE) and sub-glandular region of the stroma (SG). Arrowheads point to the glandular lumen (GL). St: stroma. Representative images shown, n=3 biopsies per group.

### *KISS1* and *KISS1R* mRNA levels decline upon decidualization of human endometrial stromal cells

The high levels of KP and KISS1R protein in stromal cells in the mid-secretory endometrium suggest a role in stromal cell decidualization. To begin studying this potential role, human endometrial stromal cells (HESCs) were decidualized *in vitro* for six days and successful decidualization was confirmed by the increase in mRNA levels of the following genes: *FOXO1, IGFBP1*, and *PRL* (**Fig. 2A-C**). FOXO1 protein levels also increased (**Fig. 2D)**, as did mRNA levels of *DCN, LEFTY2* and *WNT4* (**Fig. E-G)**. Quantitation of *KISS1* and *KISS1R* mRNA expression by qPCR on D1, D2, D4, and D6 revealed that as HESC decidualization progressed, *KISS1* mRNA levels decreased rapidly to barely detectable levels by the sixth day (**Fig. 2H**). For *KISS1R*, the response was variable (**Fig. 2I and J**). At no time was an increase in *KISS1R* mRNA levels detected during HESC decidualization. Overall, these results suggest that KP/KISS1R signaling diminishes over the course of decidualization. We therefore hypothesized that HESC decidualization requires the downregulation of the KP/KISS1R signaling system.

**Fig. 2.**
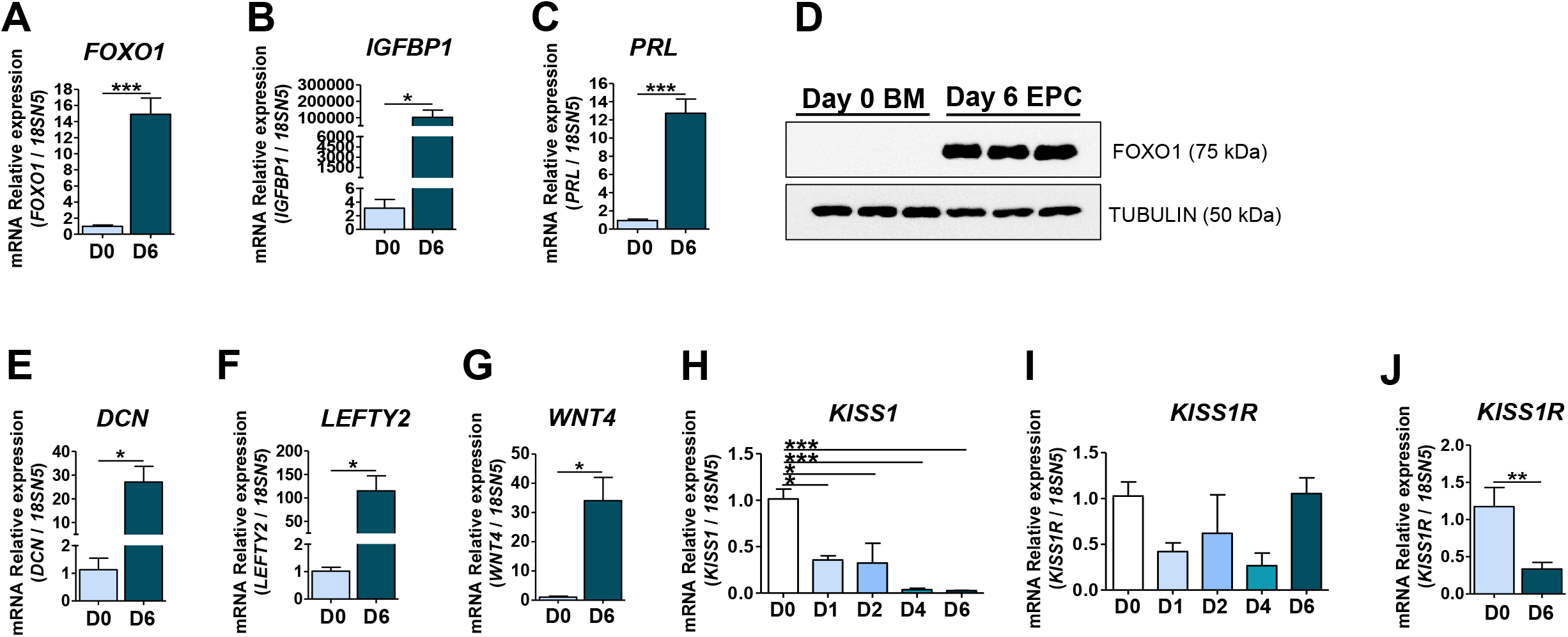
Primary ESCs undergo efficient decidualization *in* vitro and both *KISS1* and *KISS1R* mRNA levels decline with the progression HESC decidualization. Quantification of mRNA levels of hallmark molecular markers of decidualization **(A)** *FOXO1*, **(B)** *IGFBP1*, **(C)** *PRL* and (**D**) representative western blot analysis of endometrial FOXO1 protein levels with tubulin serving as the housekeeping control before and after six days of decidualization (see **Fig. S2** for additional blots). Quantification of mRNA levels of known regulators of ESC decidualization (**E**) *DCN*, (**F**) *LEFTY2*, and (**G**) *WNT4* before and after six days of decidualization. qPCR analysis of mRNA expression of (**H**) *KISS1* and (**I and J**) *KISS1R* before and after one, two, four, and six days of decidualization. Statistics (biological replicates: n=3-6 biopsies and technical replicates: n=3-6 for each group): student’s unpaired two-tailed t-test and one-way ANOVA with Bonferroni Multiple Comparison post hoc test. * p<0.05; ** p<0.01; and *** p<0.001. D0: day zero; D1: day one; D2: day two; D4: day four; D6: day six of decidualization; BM: basal medium; and EPC: decidualization medium.

### Expression of *KISS1R* in decidualized HESCs reverses the decidualized ESC phenotype

To test the hypothesis that HESC decidualization requires the downregulation of the KP/KISS1R signaling system, on the fourth day of *in vitro* decidualization, decidualizing HESCs were transfected with either a human *KISS1R* expression plasmid (containing the *KISS1R* open reading frame) or an empty expression vector (control) for two days. The goal of this study was to upregulate KP/KISS1R signaling immediately following the endogenous downregulation of the signaling system in decidualizing HESCs and determine the impact of persistent KP/KISS1R signaling on HESC decidualization. The results showed that decidualizing HESCs were efficiently transfected with exogenous *KISS1R* (**Fig. 3A**), and transfection led to an increase in the gene expression of the *KISS1R* ligand, *KISS1* (**Fig. 3B**). Furthermore, expression of *KISS1R* reversed the decidualizing HESC epithelioid phenotype based on the reduced expression of the epithelial markers *DESMIN, KRT7*, and *KRT18* (Kalluri *et al*., 2009) (**Fig. 3C**). The reduction in *KRT7* mRNA expression (**Fig. 3C**) was also observed at the protein level where compared to empty vector control (**Fig. 3D1-3**), cytokeratin 7 expression was reduced in the nuclei of decidualized HESCs transfected with *KISS1R* (**Fig. 3D4-6**). Interestingly, in HESCs, cytokeratin 7 is largely nuclear and not cytosolic (Hobbs *et al*., 2016). These findings further support the conclusion that KISS1R negatively regulates HESC decidualization by reversing the ESC decidual phenotype.

**Fig. 3.**
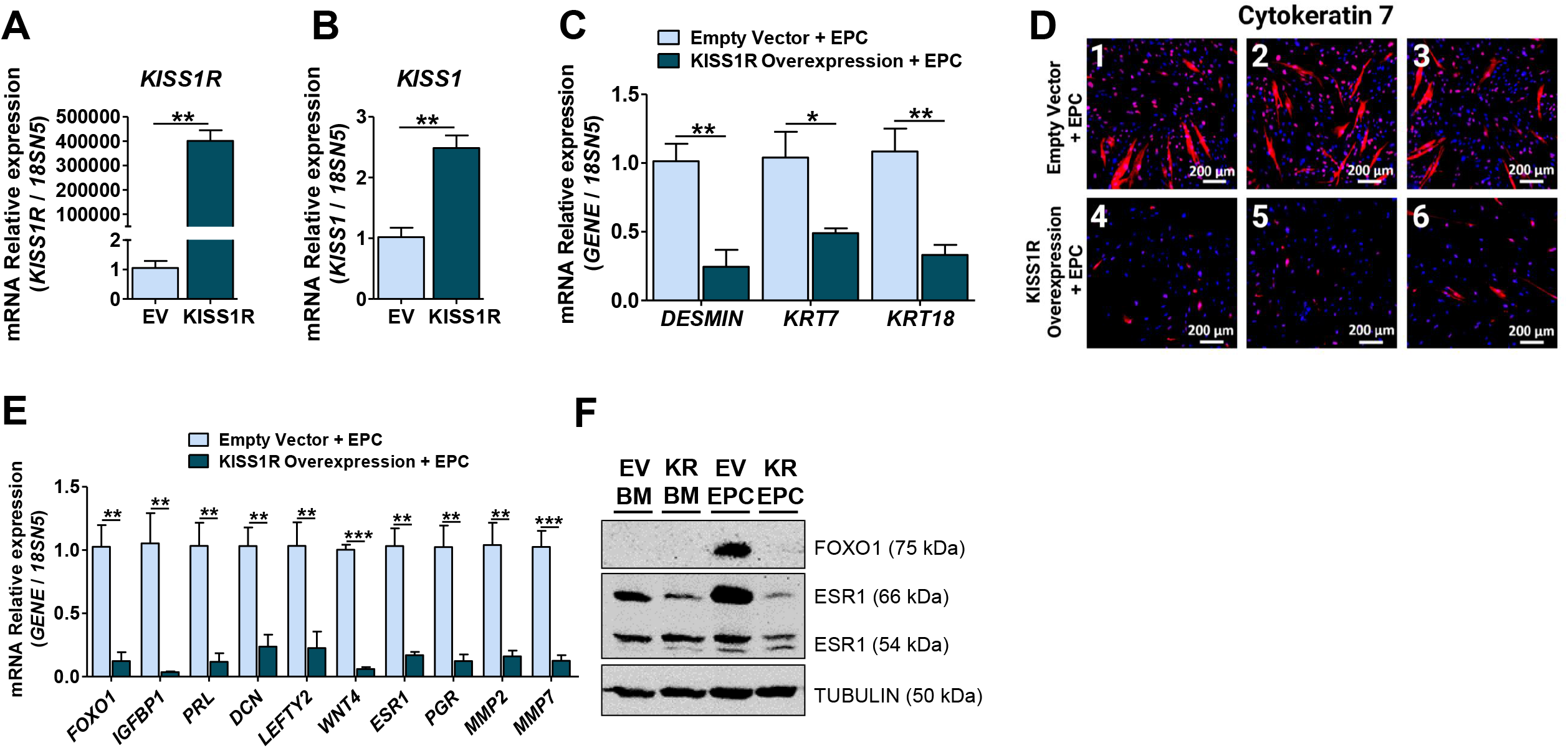
Expression of *KISS1R* in decidualizing primary ESCs reverses the decidualized ESC phenotype and decreases the expression of target markers of decidualization, ESR1 signaling, and ECM. Quantification of mRNA and protein levels in empty vector control and KISS1R expressing primary HESCs following six days of *in vitro* decidualization. qPCR analysis of (**A)** *KISS1*, (**B**) *KISS1R* and phenotypic markers of decidualization (**C**) *DESMIN, KRT7*, and *KRT18* mRNA levels. Representative images of the immunofluorescence analysis of cytokeratin 7 protein levels in empty vector control (**D1-3**) and KISS1R (**D4-6**) expressing primary HESCs. (**E**) qPCR analysis of decidualization and ESR1 signaling markers *FOXO1, IGFBP1, PRL, DCN, LEFTY2, WNT4, ESR1, PGR, MMP2*, and *MMP7* mRNA levels. Representative western blot analysis of endometrial FOXO1 and ESR1 protein levels with tubulin serving as the housekeeping control before and after six days of decidualization in empty vector and *KISS1R* expressing HESCs (see **Fig. S3** for additional blots). Statistics (biological replicates: n=3-5 biopsies and technical replicates: n=3-6 for each group): student’s unpaired two-tailed t-test. * p<0.05; ** p<0.01; and *** p<0.001. EV: Empty vector control; KISS1R or KR: KISS1R expression; BM: basal medium; and EPC: decidualization medium.

### Expression of *KISS1R* in decidualizing HESCs downregulates the expression of target markers of decidualization, ESR1 signaling, and ECM remodeling

To further test the hypothesis that HESC decidualization requires the downregulation of the KP/KISS1R signaling system, other markers of decidualization, ESR1 signaling, and ECM remodeling were analyzed in decidualizing HESCs expressing exogenous *KISS1R*. This analysis revealed that *FOXO1, IGFBP1, PRL, DCN, LEFTY2, WNT*4, *ESR1, PGR, MMP2* and *MMP7* mRNA expression levels were strongly downregulated in KISS1R-expressing cells compared to empty vector controls (**Fig. 3E**). Next, we evaluated the protein expression levels of FOXO1 and ESR1 in HESCs expressing the empty vector or *KISS1R* under basal (BM) and decidualization (EPC) conditions. This revealed that FOXO1 was only expressed under decidualization conditions when the HESCs were transfected with the empty vector, and not when they were transfected with *KISS1R* plasmid (**Fig. 3F**). Similarly, in the absence of exogenous KISS1R, the 66 and 54 kDa ESR1 isoforms are more robustly expressed under decidualizing conditions (**Fig. 3F**) but in the presence of KISS1R, expression is strikingly reduced, under both basal and decidualizing conditions (**Fig. 3F**). Taken together, these results strongly suggest that *KISS1R* negatively regulates HESC decidualization by reducing ESR1-66 kDa and ESR1-54 kDa levels and inhibiting downstream targets of decidualization.

### Expression of human *ESR1-46* in decidualized HESCs inhibits ESR1-66 and ESR1-54 signaling and reduces ESC decidualization

Since we observed that reduced ESR1-54 and ESR1-66 levels are associated with reduced decidualization in KISS1R-expressing cells (**Fig. 3F**), we hypothesized that by expressing exogenous ESR1-46 in HESCs undergoing decidualization, we could phenocopy the KISS1R expression effect. We also hypothesized that by co-expressing ESR1-66 with ESR1-46, any negative effect on decidualization would be rescued through excess exogenous ESR1-66 present in the cells. To test these hypotheses, HESCs were transfected with either empty vector control, h*ESR1-*46, or h*ESR1-*66 + h*ESR1*-46 (h*ESR1-*66/46) expression constructs under basal and decidualizing conditions. The results showed that HESCs were successfully transfected with exogenous *hESR1-* 46 and h*ESR1-*66/46 constructs based on the increased *ESR1* mRNA expression (**Fig. 4A**). The results also clearly showed that expression of h*ESR1*-46 blocked *PGR, FOXO1, IGFBP1* and *PRL* mRNA expression in cells cultured in EPC decidualization medium (**Fig. 4B** and **C**). However, when h*ESR1-66* and h*ESR1*-44 were co-transfected, this was not observed (**Fig. 4B** and **C**).

**Fig. 4.**
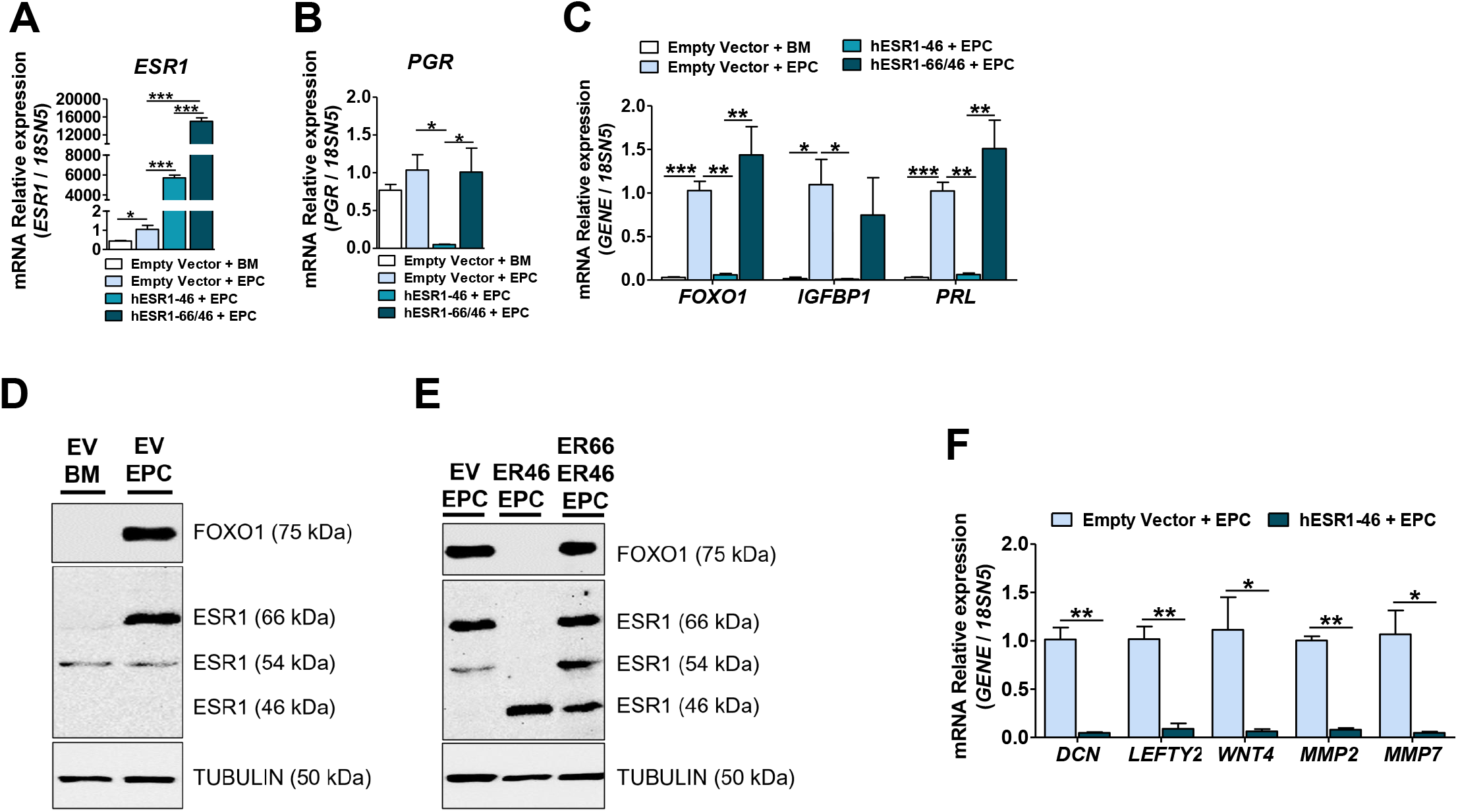
Expression of human ESR1-46 in decidualizing primary ESCs reduces ESR1-66 and ESR1-54 protein levels and thereby inhibits decidualization. Quantification of mRNA and protein levels in empty vector control, h*ESR1*-46 and h*ESR1*-66/46 expressing primary ESCs following six days of *in vitro* decidualization. qPCR analysis of (**A)** *ESR1*, (**B**) *PGR* and hallmark markers of decidualization (**C**) *FOXO1, IGFBP1*, and *PRL* mRNA levels. Representative western blot analyses of endometrial (**D and E**) FOXO1 and ESR1 protein levels with TUBULIN serving as the housekeeping control (**D**) before and after six days of decidualization in empty vector controls and (**E**) after six days of decidualization in empty vector and *hESR1-*46, and *hESR1-66/46* expressing ESCs (see **Fig. S4** for additional blots). (**F**) qPCR analysis of decidualization and ESR1 signaling markers *DCN, LEFTY2, WNT4, MMP2*, and *MMP7* mRNA levels. Statistics (biological replicates: n=3-5 biopsies and technical replicates: n=3-6 for each group): student’s unpaired two-tailed t-test and one-way ANOVA with Bonferroni Multiple Comparison post hoc test. * p<0.05; ** p<0.01; and *** p<0.001. AF-1: activation function-1 domain; DBD: DNA binding domain; H: hinge region; LBD: ligand binding domain; AF-2: activation function-2 domain; EV: Empty vector control; ER46: *hESR1*-46 expression; ER66 and ER46: *hESR1-*66/46 expression; BM: basal medium; and EPC: decidualization medium.

Next, protein analysis of HESCs cultured in EPC decidualization medium showed increased expression of the decidualization marker FOXO1 as well as ESR1-66 (**Fig. 4D**). This was identical to HESCs transfected with the empty expression vector and cultured in decidualization medium (EPC) (**Fig. 4E**). However, in cells expressing exogenous h*ESR1-*46, expression of FOXO1, ESR1-66, and ESR-1-54 proteins were ablated (**Fig. 4E**). In contrast, HESCs cultured in EPC and exogenously expressing empty vector or h*ESR1*-66/46 exhibited robust levels of FOXO1, ESR1-66, ESR1-46, and ESR1-54 protein (**Fig. 4F**).

Lastly, we quantified the expression of other downstream targets of decidualization and ESR1 signaling. Expression of exogenous h*ESR1*-46 in decidualizing HESCs resulted in the reduction of *DCN, LEFTY2, WNT4, MMP2*, and *MMP7* mRNA levels compared empty vector controls (**Fig. 4F**). Taken together, these results reveal ESR1-46 expression in decidualizing HESCs inhibits ESR1-66 and ESR1-54 protein levels leading to the downregulation of molecular targets of decidualization and ESR1 signaling, thereby diminishing ESC decidualization. Furthermore, these findings support the idea that KP/KISS1R signaling negatively regulates HESC decidualization by reducing ESR1-66 and ESR1-54 expression levels and thereby transcriptional activity.

## DISCUSSION

In women, endometrial stromal cell (ESC) decidualization begins in the mid-secretory endometrium and is localized to the luminal-distal region of the functionalis among ESCs surrounding the spiral arteries (Owusu-Akyaw *et al*., 2019; Ramathal *et al*., 2010). During this period, at the luminal-proximal end of the functionalis, an implanting embryo encounters non-decidualized ESCs which facilitate rapid stromal invasion and successful implantation. While this is occurring, the decidualization reaction spreads beyond the perivascular regions and reaches the implanted embryo restricting further invasion of the endometrium while providing nourishment and protection. We hypothesize, based on the data presented, that in order to spatially restrict decidualization to the luminal distal region of the functionalis at the start of the implantation process, decidualization must be inhibited among ESCs surrounding the implanting embryo and this is achieved through endometrial KP/KISS1R signaling. However, as our data showed, *KISS1* and *KISS1R* expression is downregulated in decidualized ESCs; thus, we suggest, following successful embryo implantation, *KISS1* and *KISS1R* expression is downregulated in ESCs surrounding the embryo and this allows them to undergo decidualization.

Our finding that KISS1R negatively regulates human ESC decidualization is in opposition to a study in mice that reported endometrial KISS1 and KISS1R are required for decidualization (Zhang *et al*., 2014). Using a mouse ESC culture model, the authors reported that *Kiss1* and *Kiss1r* mRNA expression increased with the progression of ESC decidualization, while siRNA-downregulation of *Kiss1* blocked ESC decidualization (Zhang, et al., 2014). By contrast, studies from our lab revealed that the uteri of *Kiss1* and *Kiss1r* null mice successfully underwent experimentally-induced decidualization (Calder, et al., 2014). These findings suggest that uterine KP/KISS1R signaling does not regulate ESC decidualization, or that it negatively regulates decidualization and therefore in its absence decidualization is observed. While further studies are required to understand whether KP/KISS1R signaling negatively regulates mouse ESC decidualization, our current findings show that it negatively regulates human ESC decidualization.

Studies over the years have revealed the existence of several ESR1 isoforms, that are cell- and tissue-specific in expression. While much progress has been made in understanding the functions of these isoforms, much still remains to be understood. In support of our findings, a recent report has identified the expression of the truncated splice variant of the full length ESR1-66 isoform, ESR1-46, in the nucleus and cytoplasm of the human endometrium during the proliferative and secretory phase and within first trimester decidual tissues (Gibson *et al*., 2020). Studies have revealed that ESR1-46 antagonizes the transcriptional activity of ESR1-66 by acting as a dominant negative repressor and an effective inhibitor of ESR1-66 (Chantalat, et al., 2016; Flouriot, et al., 2000; Penot, et al., 2005). In our studies, expression of h*ESR1*-46 completely abolished ESR1-66 and ESR1-54 protein levels and drastically reduced targets of decidualization and ECM remodeling comparable to that seen in HESCs expressing exogenous *KISS1R*. Furthermore, this finding is consistent with the notion that KISS1R negatively regulates HESC decidualization by reducing ESR1 expression and transcriptional activity.

In conclusion, we our data support the notion that endometrial KISS1R positively regulates embryo implantation by negatively regulating *ESR1* expression. This ensures that ESR1 transcriptional activity is kept in check at time of embryo implantation to allow the epithelium to acquire a receptive state and delay the period prior to stromal fibroblast decidualization as embryos are unable to implant in decidualized cells. After the embryo has implanted, KISS1R signaling decreases, allowing stromal cells surrounding the embryo to decidualize, thereby preventing the embryo from undergoing further and aberrant invasion of the stroma while at the same time nourishing and protecting the developing embryo until the functional placenta forms.

## Non-standard abbreviations

db-cAMP: N6,2’-O-dibutyryladenosine 3’:5’-cyclic monophosphate
*DCN*: Decorin
ESC: Endometrial stromal cell
ESR1: Estrogen receptor alpha, ERα
*FOXO1*: Forkhead box protein O1
HESC: Human endometrial stromal cell
*IGFBP1*: Insulin-like growth factor binding protein 1
*KISS1*: Kisspeptin 1 gene
*KISS1R*: Kisspeptin receptor
KP: Kisspeptin protein
*KRT7*: Cytokeratin 7
*KRT18*: Cytokeratin 18
LEFTY2: Left-right determination factor 2
LH: Luteinizing hormone
MPA: Medroxyprogesterone acetate
*MMP2*: Matrix metallopeptidase 2
*MMP7*: Matrix metallopeptidase 7
*PGR*: Progesterone receptor gene
PR: Progesterone receptor protein
*PRL*: Prolactin
*WNT4*: Wingless-type MMTV integration site family member 4

## ACKNOWLEDGEMENTS

We thank Dr. Gilles Flouriot (University of Rennes, Inserm, EHESP, Institut de Recherche en Santé, Environnement et Travail, UMR_S 1085, F-35000, Rennes, France) for his kind gifts of the human *ESR1-*66 and *ESR1*-46 DNA constructs. JS was supported by the NIH/NIEHS Training Grant ES007148, and additional funding for the study was obtained from the Busch Biomedical Grant (to AVB).

## AUTHOR CONTRIBUTIONS

A. V. Babwah developed the research study and all authors contributed to the analysis of the data and preparation of the manuscript.

## DATA AVAILABILITY

All data generated or analyzed during this study are included in this published article.

