## Supplementary material for "Human endometrial KISS1R inhibits stromal cell decidualization in a manner associated with a reduction in ESR1 levels": Suplemental Figure Legends

### SUPPLEMENTAL DATA

**Supplemental Fig. 1.** Diagram illustrating the structure and domains within the conserved regions A-F of the estrogen receptor alpha (ESR1) isoforms: full length ESR1-66 and truncated isoforms ESR1-54 lacking exon 4 and the following residues 264 to 302 (denoted by ▲4) within the hinge region and ESR1-46 lacking an N-terminal AF-1 domain.

**Supplemental Fig. 2. Primary ESCs undergo efficient decidualization *in vitro*.** Western blot analysis of FOXO1 protein levels with tubulin serving as the housekeeping control before and after six days of decidualization. (**A-C**) Biopsy ID#: UC742, (**D-F**) biopsy ID#: UC744, and (**G and H**) biopsy ID#: UC7481. BM: basal medium; and EPC: decidualization medium.

**Supplemental Fig. 3. Expression of *KISS1R* in decidualizing primary ESCs decreases the protein expression of ESR1-66, ESR1-54 and FOXO1.** Western blot analysis of FOXO1 and ESR1 protein levels with tubulin serving as the housekeeping control before and after six days of decidualization in empty vector and *KISS1R* expressing ESCs. (**A-D**) Biopsy ID#: UC742, (**E-G**) biopsy ID#: UC744, and (**H-K**) biopsy ID#: UC7481. EV: Empty vector control; KISS1R or KR: KISS1R expression; BM: basal medium; and EPC: decidualization medium.

**Supplemental Fig. 4. Expression of *KISS1R* in decidualizing primary ESCs decreases the protein expression of ESR1-66, ESR1-54 and FOXO1.** Western blot analysis of FOXO1 and ESR1 protein levels with tubulin serving as the housekeeping control (**A-D**) Biopsy ID#: UC742, (**I-K**) biopsy ID#: UC744, and (**O-R**) biopsy ID#: UC7481 before and after six days of decidualization in empty vector controls and (**E-H**) Biopsy ID#: UC742, (**L-N**) biopsy ID#: UC744, and (**S-V**) biopsy ID#: UC7481 after six days of decidualization in empty vector and *hESR1*-46, and *hESR1*-66/46 expressing ESCs. EV: Empty vector control; ER46: *hESR1*-46 expression; ER66 and ER46: *hESR1*-66/46 expression; BM: basal medium; and EPC: decidualization medium.
