## Supplementary figures and images for "Human endometrial KISS1R inhibits stromal cell decidualization in a manner associated with a reduction in ESR1 levels"

### Supplemental Figures

#### Supplemental Figure 1

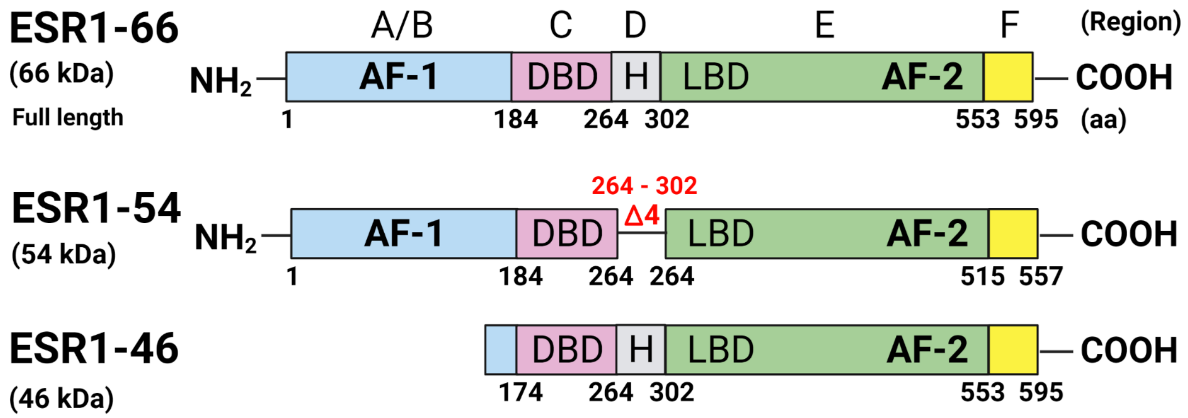

#### Supplemental Figure 2

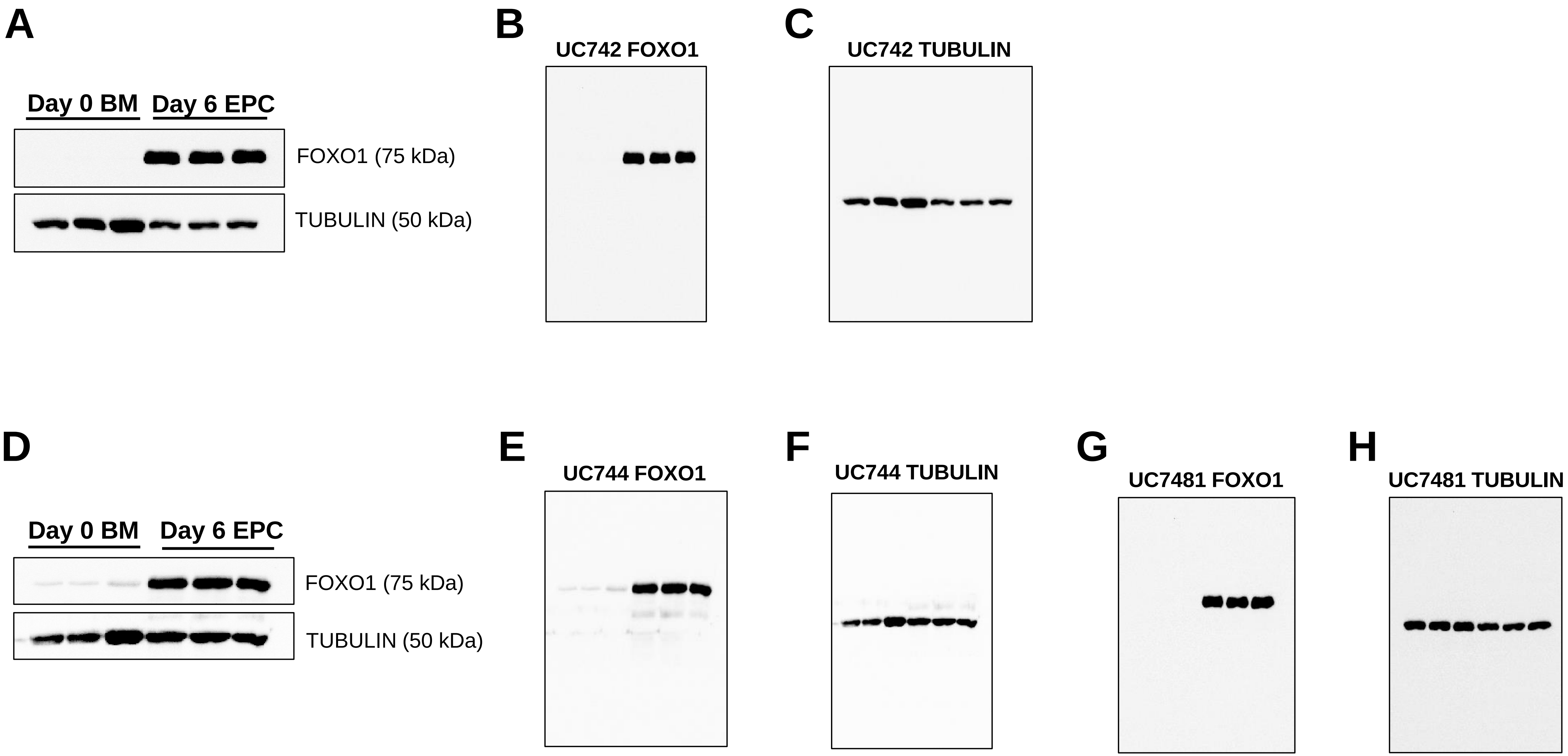

#### Supplemental Figure 3

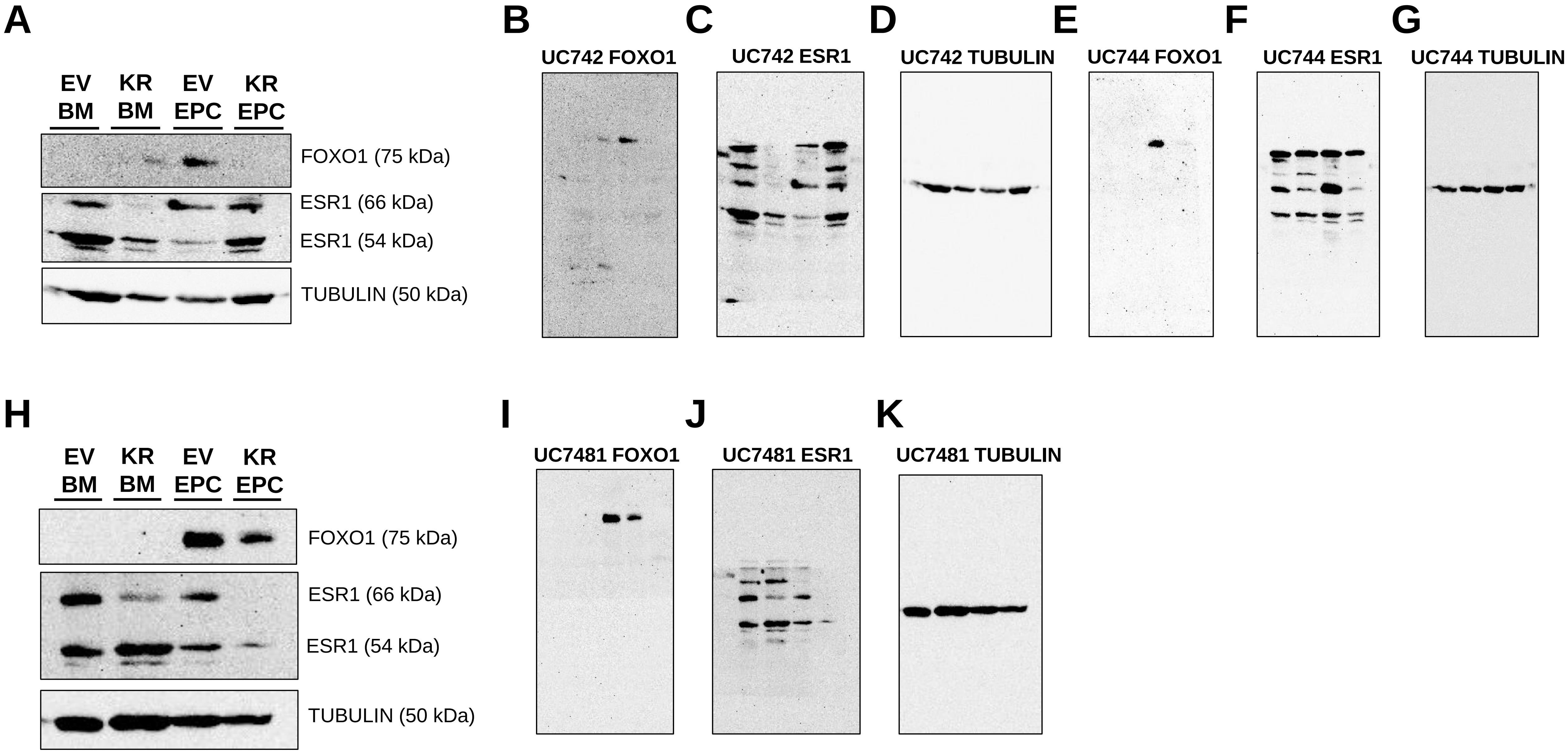

#### Supplemental Figure 4

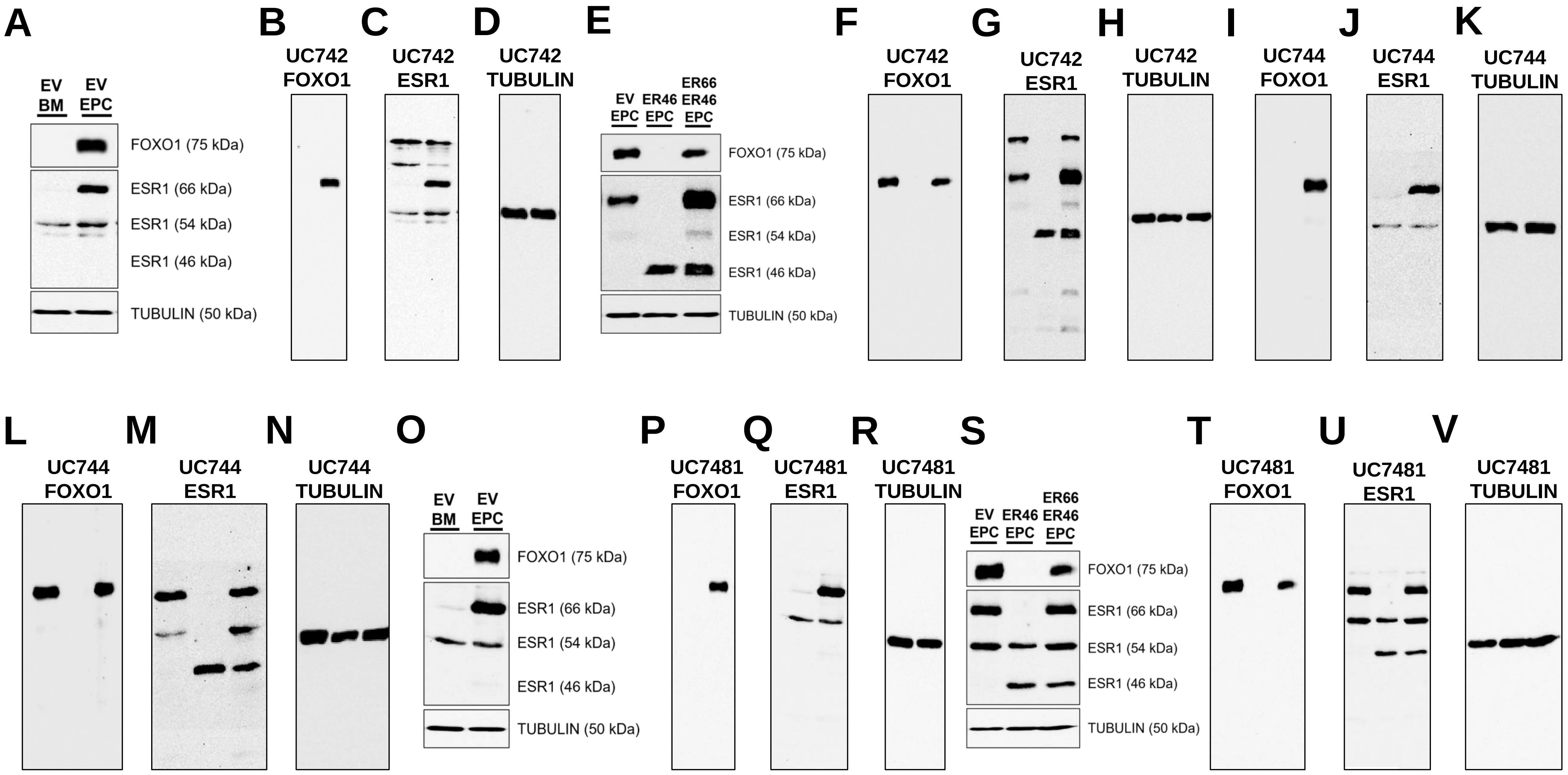
